# Dissociation of immediate and delayed effects of emotional arousal on episodic memory

**DOI:** 10.1101/242511

**Authors:** Dirk Schümann, Janine Bayer, Deborah Talmi, Tobias Sommer

## Abstract

Emotionally arousing events are usually better remembered than neutral ones. This phenomenon is in humans mostly studied by presenting mixed lists of neutral and emotional items. An emotional enhancement of memory is observed in these studies often already immediately after encoding and increases with longer delays and consolidation. A large body of animal research showed that the more efficient consolidation of emotionally arousing events is based on an activation of the central noradrenergic system and the amygdala (Modulation Hypothesis; Roozendaal & McGaugh, 2011). The immediately superior recognition of emotional items is attributed primarily to their attraction of attention during encoding which is also thought to be based on the amygdala and the central noradrenergic system. To investigate whether the amygdala and noradrenergic system support memory encoding and consolidation via shared neural substrates and processes a large sample of participants (n = 690) encoded neutral and arousing pictures. Their memory was tested immediately and after a consolidation delay. In addition, they were genotyped in two relevant polymorphisms (α_2B_-adrenergic receptor and serotonin transporter). Memory for negative and positive emotional pictures was enhanced at both time points where these enhancements were correlated (immediate r = 0.60 and delayed test r = 0.46). Critically, the effects of emotional arousal on encoding and consolidation correlated only very low (negative r = 0.14 and positive r = 0.03 pictures) suggesting partly distinct underlying processes consistent with a functional heterogeneity of the central noradrenergic system. No effect of genotype on either effect was observed.

## 1 Introduction

### 1.1 Enhanced consolidation of emotionally arousing events

Emotionally arousing events are usually remembered better than neutral ones, a phenomenon called the emotional enhancement of memory (EEM). A large body of animal data have shown that the EEM is caused by a rise in peripheral stress hormone levels induced by an emotionally arousing event which activates the central noradrenergic (NOR) system, based in the locus coeruleus (LC) via ascending fibers (Modulation Hypothesis; McIntyre, McGaugh, & Williams, 2012). The LC projects to nearly all brain areas including the basolateral amygdala (BLA) which in turn modulates memory formation in the hippocampus. Both tonic and phasic activations of the BLA result in an EEM; for example, a brief electrical stimulation of the BLA also produces a stimulus-specific EEM, observed in the altered exploration of neutral objects that preceded the BLA stimulation (Bass, Partain, & Manns, 2012; Bass, Nizam, Partain, Wang, & Manns, 2014). Critically, the EEM that results from an increase in stress hormone levels and BLA activations is not observed immediately but only after a consolidation delay - consistent with the *Emotional Synaptic Tagging Hypothesis* (Bergado, Lucas, & Richter-Levin, 2011; McReynolds & McIntyre, 2012).

### 1.2 Studies in humans on the emotional enhancement of memories

The majority of studies on the EEM in humans, in particular nearly all fMRI experiments, investigate the effect of emotional arousal on memory formation by presenting mixed lists of emotional and neutral items. Better memory for emotional items is thought to be a laboratory measure of the EEM. At the neural level the EEM is associated with greater activity in the amygdala, hippocampus, and parahippocampus, in addition to visual, prefrontal, and parietal areas (Dolcos, Denkova, & Dolcos, 2012; LaBar & Cabeza, 2006; Murty, Ritchey, Adcock, & LaBar, 2010). Thus, the behavioral and neural results appear similar to what has been described in the animal literature on the EEM.

Yet there are critical differences between human fMRI studies of the EEM and animal studies. The rapid stimulus-specific EEM in fMRI-studies is comparable to stimulus-specific EEM elicited by electrical BLA stimulation or by a direct NOR infusions into the BLA (shown to enhances memory for previously-explored objects, Barsegyan, McGaugh, & Roozendaal, 2014; Bass et al., 2012), but it cannot be caused by a relatively slow, sluggish systemic increase in stress hormone levels, which characterizes the majority of animal research on EEM. Human fMRI studies also contrast with stimulus-specific animal experiments. While in these experiments the target stimuli are neutral objects, human fMRI studies compare neutral stimuli to stimuli that themselves evoke emotional arousal as a result of their semantic evaluation, and are hence already differentially encoded. Most importantly, in these experiments an EEM is often observed also when memory is tested almost immediately after encoding (Murty et al., 2010; Talmi & McGarry, 2012), an effect that cannot be caused by more efficient consolidation (Talmi, 2013).

### 1.3 Immediately enhanced emotional memories

Immediate EEM has been explained in cognitive psychology by three characteristics of emotional stimuli. In particular, emotional stimuli are semantically more closely related, they are more distinct and attract more selective attention (Sommer, Glascher, Moritz, & Buchel, 2008; Talmi, 2013). Memory undoubtedly benefits from the attention emotionally-arousing stimuli capture during encoding. Enhanced attention to such stimuli, as reflected, for instance, in more fixations, results in deeper processing during encoding and better immediate memory (Bradley, Houbova, Miccoli, Costa, & Lang, 2011; Kang, Wang, Surina, & Lü, 2014; Miendlarzewska, van Elswijk, Cannistraci, & van Ee, 2013; Sharot & Phelps, 2004; Talmi, Anderson, Riggs, Caplan, & Moscovitch, 2008). Moreover, enhanced attention to emotional stimuli during encoding is also consistent with the more frequent experience of recollection during the recognition of emotional items (Kensinger, Clarke, & Corkin, 2003; Kensinger & Corkin, 2003). Critically, the enhanced immediate recollection of emotional items correlates with amygdala activity during encoding (Kensinger, Addis, & Atapattu, 2011) implicating an encoding related effect that is independent from the more liberal response bias which is induced by their higher semantic relatedness and influences the recognition of emotional stimuli (Dougal and Rotello 2007). While increased attention can increase episodic memory in recognition as well as in free recall tests, the other characteristics of emotional stimuli known to increase immediate memory, i.e. their higher semantic relatedness and distinctiveness, might primarily enhance recall but not recognition accuracy. Taken together, the EEM observed in recognition tests after a study-test delay too short for consolidation probably mainly reflects increased attention to emotionally-arousing stimuli during encoding (Hamann, 2001; Todd, Palombo, Levine, & Anderson, 2011). Interestingly, studies that elevated arousal only after initial processing of neutral stimuli and observed no immediate but only enhanced delayed memory have suggested a partial independence of arousal-induced processes on encoding and consolidation (Bass et al., 2012; Schwarze, Bingel, & Sommer, 2012).

### 1.4 Aim of the current study: Do the effects of emotional arousal on consolidation and immediate memory rely on the same neural substrate?

The current study aimed to further characterize the relationship between the effect of emotional arousal on immediate and delayed recognition. In particular, we aimed to find out whether both effects rely on the same neural substrate. The current experimental approach follows the rationale that if both effects stem from the same underlying neural circuits, their magnitudes should be correlated across participants. Therefore, we administered an emotional memory paradigm with immediate and delayed recognition tests to a large sample of participants and correlated the immediate and delayed EEM. In order to be maximally sensitive to differences in the effects of arousal the EEMs were assessed not only as differences in recognition accuracy (corrected hit rate and d-prime) but also in terms of response bias, discriminability (bias-corrected accuracy), and the separate contributions of arousal to familiarity and recollection (White, Kapucu, Bruno, Rotello, & Ratcliff, 2014; Yonelinas, 1994). In order to illuminate potential reasons for differences in immediate and delayed EEMs across participants they were characterized by several neuropsychological tests and relevant questionnaires.

In addition, volunteers were genotyped in two polymorphisms in the genes coding for the α_2B_-noradrenergic receptor and the serotonin transporter in order to associate the immediate and delayed EEM with the different genotypes. The first polymorphism reduces the α_2B_-noradrenergic receptor functionality and is in a complete linkage disequilibrium with a polymorphism that results in less transcription of the α2B-noradrenergic receptor (Crassous et al., 2010; Nguyen, Kassimatis, & Lymperopoulos, 2011; Salim, Desai, Taneja, & Eikenburg, 2009; Small, Brown, Forbes, & Liggett, 2001). Carriers of the allele are expected therefore to have fewer and less functional α_2B_-noradrenergic receptors. This variant has been associated with the greater vividness of processing emotional stimuli, the attraction of selective attention and the immediate EEM in free recall (de Quervain et al., 2007; Rasch et al., 2009; Todd et al., 2013, 2015). The polymorphism in the gene coding for the serotonin transporter has been associated with attentional bias to negative stimuli and greater amygdala reactivity (Bevilacqua & Goldman, 2011; Canli, Ferri, & Duman, 2009; Munafò, Brown, & Hariri, 2008). The rationale of this complementary experimental approach was that an association of both the immediate and delayed EEM with these polymorphisms would suggest shared underlying neural substrates.

## 2 Methods

### 2.1 Participants

The present emotional memory data were taken from a sample of 690 young healthy adults, who took part in a larger test and questionnaire battery to phenotype participants with respect to a variety of cognitive and affective characteristics. 44 participants were excluded due to incomplete data or below chance performance, resulting in a sample of n = 646 (464 females, age range 18–36 y, mean 24.4 y).

### 2.2 Emotional memory paradigm

Stimulus material were 240 pictures of different valence and arousal levels (80 negative, 80 positive, 80 neutral) taken from the IAPS and the internet. An independent sample (n = 52) rated valence and arousal of all pictures using respective 9-step SAM scales (Figure 1 A; Bradley & Lang, 1994). Means of arousal and valence between negative (valence: 2.05; arousal: 6.99), neutral (6.05; 3.11) and positive pictures (7.75; 3.51) were different at p < 0.0001 (except positive arousal greater than neutral p < 0.01). Note that the average arousal and valence difference between emotional and neutral items were less pronounced for positive than for negative pictures. The pictures were matched between emotional categories so that each category contained an equal number of pictures with a similar content (e.g. two interacting humans, an animal, food). Within each category pictures with similar content were for each participant individually pseudorandomly assigned to target and lures.

**Figure 1:**
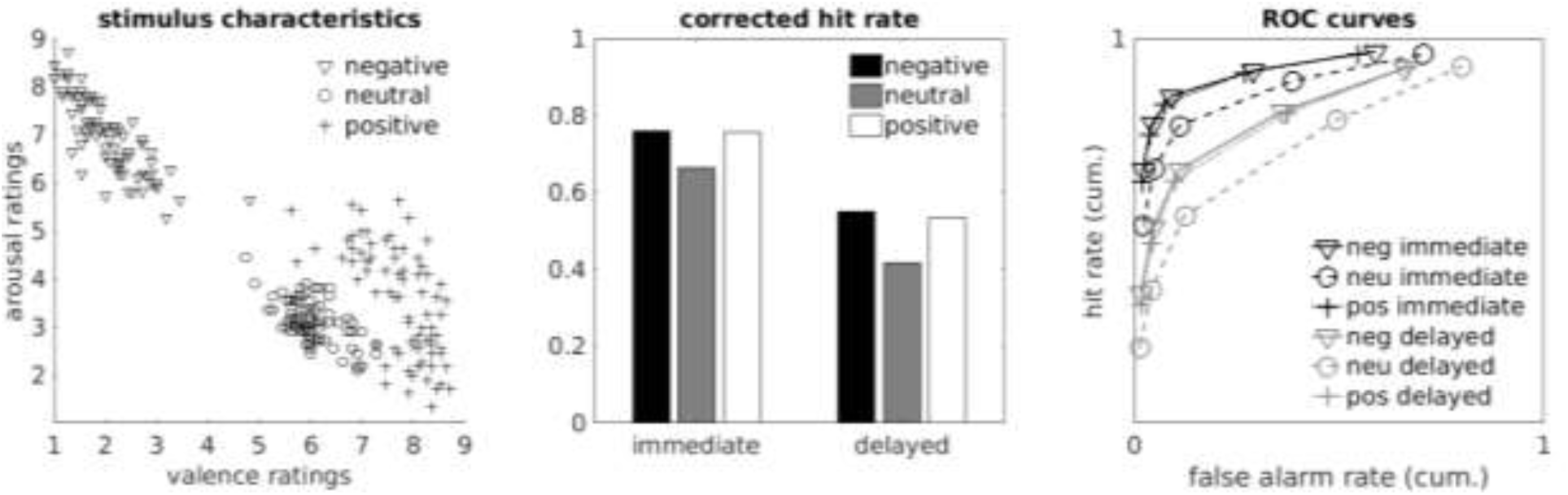
Stimuli characteristics and memory performance. A) Mean valence and arousal ratings of the used stimuli. B) Corrected hit rate in the immediate and delayed recognition test. C) ROC-curves for the immediate and delayed test.

Participants incidentally encoded the mixed list of emotionally arousing and neutral pictures (40 negative, 40 positive and 40 neutral) by rating on a 6 point Likert scale whether the picture would be suitable for a news magazine. Participants sat within normal distance of a 24” monitor and responded via keyboard arrow keys to the task presented with Presentation software (Neurobehavioral Systems). Color photographs of size 375 × 280 pixel were presented at the center of the screen on a black background for 1 s, followed by a 2-s rating interval and a 3 s active baseline task (three runs of a left/right button press in response to an arrow). 10 minutes later, filled with a divided attention task (Zimmermann & Fimm, 2007), 50% of targets (20 pictures of each emotion category) were presented again mixed with an equal number of lures and participants indicated their recognition memory without temporal constraint using a 6 point confidence scale ranging from *sure old* to *sure new* (immediate recognition test). Recognition memory for the remaining 50% of targets and lures was tested again 20 hours later with the identical procedure (delayed recognition test).

To rule out potential confounds induced by the preceding or subsequent tests in the context of the test battery, an independent replication sample of 45 participants (24 females, mean age 24 y) completed the same emotional memory paradigm.

### 2.3 Questionnaires and neuropsychological testing

Of all tests administered in the phenotyping effort, only all tests and questionnaires assessing emotional processing and related disorders and are hence of relevance for the current study will be mentioned here. The questionnaires that are related to emotional processing were cognitive emotion regulation (CERQ; Garnefski & Kraaij, 2006), psychopathological symptoms (SCL90-R; Franke, 2002), and trait anxiety (STAI-T; Laux, Glanzmann, Schaffner, & Spielberger, 1981). To further assess cognitive and affective characteristics, participants performed tests for crystallized as well as fluid intelligence/reasoning (recognition vocabulary test/visual and lexical matrices), working memory (computerized versions of forward and backward digit span and spatial span), attention (subtest *divided attention* of TAP, (Zimmermann & Fimm, 2007), memory and spatial source memory for neutral words. As mentioned above, participants were additionally genotyped for polymorphisms in the genes coding for the α_2B_-adrenoceptor and a serotonin transporter (Todd et al., 2011).

## 3 Results

### 3.1 Effects of emotional arousal on immediate and delayed memory performance

**Analysis of corrected hit rate**. The emotional memory paradigm produced an EEM in the immediate and delayed recognition test which will be termed immediate and delayed EEM respectively (Figure 1 B; Table 1). A repeated-measures GLM with *corrected hit rate* (hit rate - false alarm rate) as dependent variable (DV) and the factors *valence* (neutral vs. negative vs. positive) and *retention interval* (immediate vs. delayed) showed significant main effects as well as an interaction. Memory was higher in the immediate than in the delayed test (F_(1,645)_= 2285.02, p < 0.0001, partial η^2^ = 0.78) and differed between emotion categories (F_(2,1290)_= 347.02, p < 0.0001, partial η^2^ = 0.49). The significant interaction of time and valence (F_[2,1290]_ = 8.52, p < 0.0001) indicates stronger delayed compared to immediate EEM (post hoc, negative: t_[653]_ = 4.05, p < 0.0001; positive: t_[653]_ = 2.71, p = 0.007). Post hoc tests revealed EEM for negative and positive pictures in the immediate test (p < 0.0001, Cohens d = 0.57 and .55) and even more pronounced after 20 hours (p < 0.0001, d = 0.73 and d = 0.64). Memory for positive and negative items did not differ at either retention interval (p > .13, Bonferroni-corrected). The effect of positive and negative arousal on immediate as well as on delayed memory correlated significantly across participants (immediate EEMs: r=0.60, p<0.0001; delayed EEMs: r=0.46, p<0.0001), although the correlation was stronger for immediate EEM (z = 3.50, p < 0.001 (Diedenhofen & Musch, 2015).

**Table 1.** Memory performance in the immediate and delayed recognition tests using the various dependent variables. (Mean and standard deviation).

|  | hits | false alarms | c | d' | familiarity | recollection | AUC |
| --- | --- | --- | --- | --- | --- | --- | --- |
| immediate test |  |  |  |  |  |  |  |
| negative | 0.85 (0.12) | 0.09 (0.09) | 0.14 (0.34) | 2.45 (0.70) | 1.64 (0.73) | 0.51 (0.29) | 0.92 (0.07) |
| neutral | 0.77 (0.16) | 0.11 (0.11) | 0.22 (0.38) | 2.07 (0.70) | 1.43 (0.69) | 0.39 (0.28) | 0.87 (0.09) |
| positive | 0.83 (0.13) | 0.07 (0.08) | 0.22 (0.32) | 2.44 (0.69) | 1.63 (0.72) | 0.48 (0.30) | 0.91 (0.07) |
| delayed test |  |  |  |  |  |  |  |
| negative | 0.66 (0.17) | 0.11 (0.09) | 0.41 (0.38) | 1.70 (0.61) | 1.20 (0.56) | 0.25 (0.21) | 0.83 (0.09) |
| neutral | 0.54 (0.19) | 0.12 (0.10) | 0.55 (0.39) | 1.30 (0.60) | 0.91 (0.49) | 0.15 (0.16) | 0.75 (0.11) |
| positive | 0.63 (0.16) | 0.10 (0.09) | 0.48 (0.33) | 1.66 (0.62) | 1.16 (0.50) | 0.23 (0.19) | 0.82 (0.09) |

During encoding, participants rated “the suitability of the pictures for a news magazine” on a 6 point Likert scale. The rate of pictures rated suitable (4–6) was higher for negative (0.61) and positive (0.59) than for neutral pictures (0.48; F_[2,1286]_ = 69.98, p < 0.0001). A post-hoc test showed no difference in perceived suitability between negative and positive pictures (t_[643]_ = 1.59, p = 0.11). Pictures that received a less frequent rating might become more distinctive, which could affect memory. This possibility was examined by comparing the corrected hit rates for pictures associated with the less-and more-common rating in a repeated measures GLM with corrected hit rate as DV and factors rating frequency (less vs. more common suitability rating), valence and retention interval as factors). However, neither the main effect of rating frequency F_(1,561)_ = 0.17, p = 0.68, nor the interactions rating frequency x valence, F_(2,1122)_ = 0.11, p = 0.89, or rating frequency x valence x delay, F_(2,1122)_ = 0.71, p = 0.49.

**Analysis of additional accuracy measures**. Because it has been previously observed that the EEM in recognition tests might be based on a more liberal response bias for emotional items rather than augmented accuracy (White et al., 2014) we additionally computed d-prime, response bias (c), and -as a pure measure of accuracy (or discrimination; Dougal & Rotello, 2007) - the area under the curve (AUC) of the ROC-curves based on the confidence ratings (Table 1, Figure 1 C). Finally, to assess the contribution of recollection and familiarity to the EEMs the dual process model of recognition memory was fit to the individual data using the ROC toolbox (Koen, Barrett, Harlow, & Yonelinas, 2016). However, the memory paradigm comprised substantially fewer trials per condition (valence x delay) than recommended (20 instead of 60; Yonelinas & Parks, 2007) which partly resulted in poor model fits and implausible familiarity estimates. Participants with a familiarity estimate exceeding 4 were excluded (based on a normal distribution fitted to the estimates across all conditions), resulting in a sample size of n = 398 for familiarity and recollection analyses. Repeated-measures analyses produced similar results for AUC and recollection as for corrected hits (significant main effects and interaction). Familiarity estimates and d-prime also showed main effects, but no interaction (Table 2).

**Table 2.** Results of the valence x retention interval rmANOVAs and the correlation of the immediate (iEEM) and delayed EEM (dEEM) for negative and positive pictures using the various dependent variables. Greenhouse-Geisser correction for the violation of the iid-assumption was applied where necessary. ** p < 0.01.

|  | rmANOVA |  |  |  |  |  | Correlation |  |
| --- | --- | --- | --- | --- | --- | --- | --- | --- |
|  | retention interval |  | valence |  | ret. int.* valence |  | negative<br>iEEM x<br>dEEM | positive<br>iEEM x<br>dEEM |
|  | df | F | df | F | df | F |  |  |
| d' | 1,645 | 1,834.30** | 2,1290 | 266.08** | 2,1290 | 0.17 | 0.06 | 0.02 |
| AUC | 1,645 | 2,136.14** | 1.9,1241.5 | 394.46** | 2,1290 | 30.25** | 0.17** | 0.13** |
| familiarity | 1,397 | 428.10** | 2, 794 | 50.48** | 1.9,780.2 | 1.27 | 0.06 | 0.03 |
| recollection | 1,397 | 623.74** | 1.9,792,3 | 72.10** | 2,794 | 3.78* | 0.12* | 0.14** |
| c | 1,645 | 674.85** | 2,1290 | 50.09** | 2,1290 | 6.96** | 0.18** | 0.14** |

**Analysis of response bias**. This analysis showed a slightly different pattern. Generally, bias shifted to more conservative from immediate to delayed test. Immediately, a more liberal criterion was observed only for negative pictures. At the longer retention interval Bonferroni-corrected post hoc tests revealed significantly more conservative criteria for all valence categories, with the criterion for negative pictures still most liberal (negative < positive < neutral).

### 3.2 Correlation of the effects of emotional arousal on immediate and delayed memory

To explore our primary research question and to test whether individuals showing a large effect of emotional arousal at encoding (operationalized through immediate EEM) also exhibit a large effect on consolidation (operationalized through delayed EEM) and vice versa we correlated the immediate and delayed EEMs across participants. This relationship was significant, but rather subtle (corrected hit rate; negative EEM: r = 0.14, p < 0.0005; positive EEM: r = 0.03, p = 0.38; Figure 2 A). A similarly low correlation was obtained between the other DVs (d-prime, c, AUC, recollection, familiarity) associated with immediate and delayed EEMs (Table 2). Similar low correlations were observed in the replication sample (corrected hit rate; negative: r = −0.09, p = 0.76; positive r = 0.19, p = 0.23).

**Figure 2.**
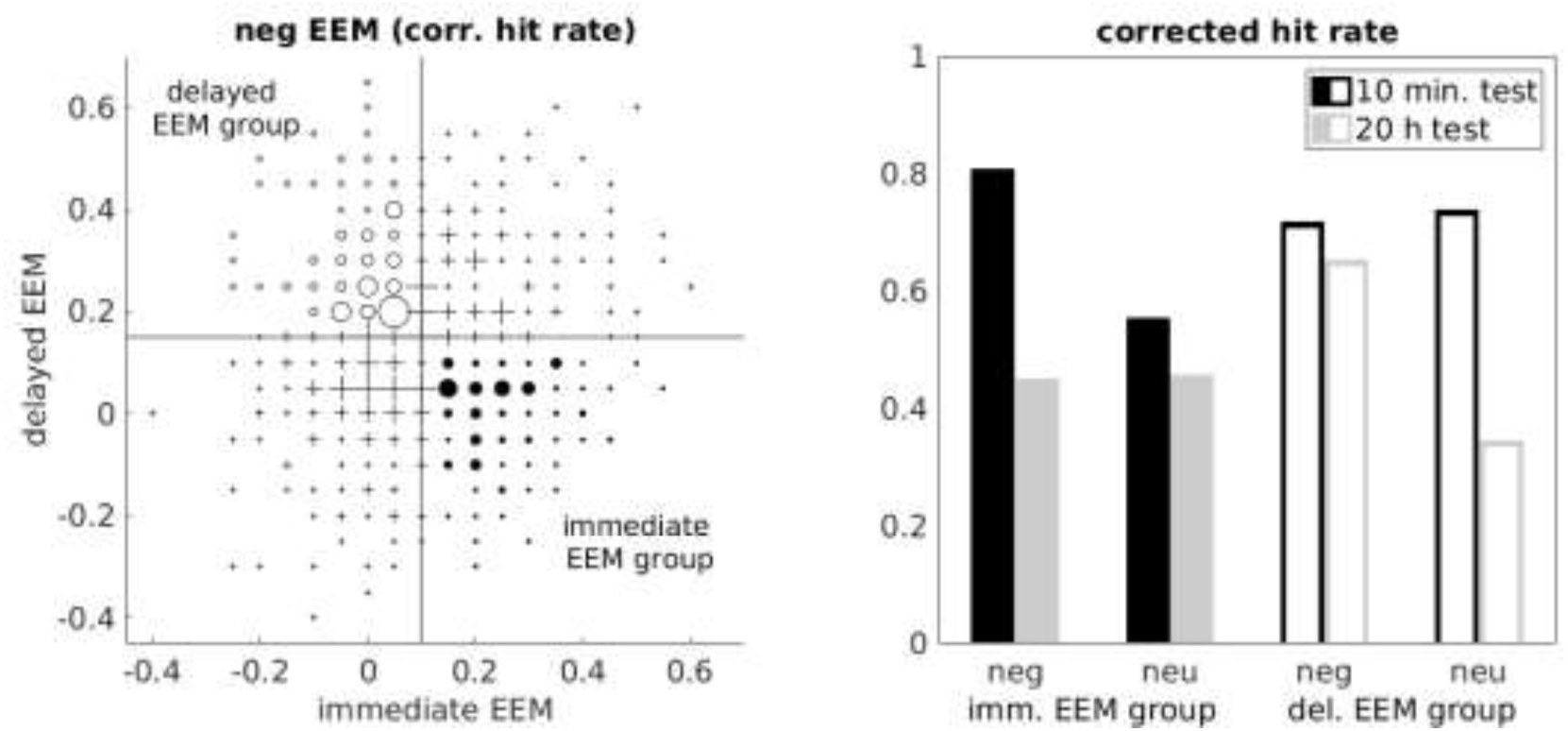
Relationship of retention interval and the negative EEM across participants. (A) Distribution of the immediate and delayed negative EEM. Horizontal and vertical lines depict median of immediate and delayed EEM respectively. Participants in the lower right corner show an above median immediate and a below median delayed EEM (immediate EEM group). Participants in the upper left corner show the opposite pattern of EEMs (delayed EEM group). The size of the data points symbolize the number of participants sharing exactly the same result. (B) Mean memory performance of the immediate and delayed negative EEM groups.

The EEM scores we correlated above are difference scores (negative minus neutral corrected hit rate) of difference scores (hits minus falls alarm rates). Differences scores have been criticized for low reliability (Gollwitzer, Christ, & Lemmer, 2014). In turn, the observed correlation of unreliable measures is lower than the true correlation (Vul, Harris, Winkielman, & Pashler, 2009). We were therefore concerned that the low correlation of the immediate and delayed EEMs might be only a result of the low reliability of the EEMs. In order to account for this confound we estimated the true correlation of the immediate and delayed EEM by correcting for their reliability. The reliability of difference scores depends on the reliabilities of their single measures as well as their correlation and the ratio of their variances (Trafimow, 2015). First, we computed the mean split-half reliabilities (Spearman-Brown corrected) for immediate and delayed hit and false alarm rates for neutral, positive and negative pictures. Therefore, for the immediate and delayed test the 20 neutral, 20 positive and 20 targets as well as the same amount of lures were across 1000 permutations randomly divided into two halves. In each permutation the correlation between the two halves of hit and false alarm rates for each valence was computed and Spearman-Brown corrected (for brevity we report here and in the remaining paragraph only the mean split-half reliabilities across valences: immediate hit rates r = 0.89 ± 0.01, delayed hit rate r = 0.80 ± 0.02, immediate false alarm rate r = 0.57 ± 0.05, delayed false alarm rate r = 0.53 ± 0.05). With these reliabilities we adjusted the observed correlations between hit and false alarm rates for all valences in order to estimate the true correlation using the formulas given in Trafimow (2015) (mean true correlation of hits and false alarm rates across valences: immediate r = −0.041 ± 0.09, delayed r = 0.20 ± 0.10). Based on the reliabilities, the true correlations and the variance ratios of hits and false alarms (mean variance ratio across valences: immediate σ_false alarms_/σ_hits_ = 0.47 ± 0.11, delayed σ_false alarms_/σ_hits_ = 0.30 ± 0.005) we computed the reliabilities of the immediate and delayed corrected hit rates for the three valences (mean reliability across valences: immediate r = 0.83 ± 0.02, delayed r = 0.73 ± 0.02). The reliabilities of the immediate (positive r = 0.66, negative r = 0.63) and delayed EEMs (positive r = 0.55, negative r = 0.52) were obtained by repeating the described procedure using true correlations between the negative, positive and neutral corrected hit rates (immediate correlations: positive and neutral r = 0.66, negative and neutral r = 0.68; delayed correlations: positive and neutral r = 0.72, negative and neutral r = 0.78) and variance ratios (immediate: σ_neutral_/σ_positive_ 1.43,σ_neutral_/σ_negative_ 1.45, delayed σ_neutral_/σ_positive_ 1.12,σ_neutral_/σ_negative_ = 1.15). Adjusting the observed correlations with these reliabilities gives true correlation estimate of r = 0.06 for the positive and r = 0.24 for the negative immediate and delayed EEMs.

In order to further explore the low correlation between the immediate and delayed EEMs, we used the corrected hit rate to identify two extreme groups by median splits of the immediate and delayed EEM (Figure 2 A). Since the animal work leading to the *Modulation* and *Emotional Synaptic Tagging Hypotheses* is based on negative events (Roozendaal & McGaugh, 2011), we focused the following analyses on the negative EEMs. The ‘immediate negative EEM’ group was defined as participants who exhibited an immediate negative EEM above the median but a delayed negative EEM below the median. In that group negative emotional arousal had a large effect on encoding (Figure 2B) but not on consolidation, because participants forgot disproportionally many negative pictures overnight. The delayed negative EEM group was defined as participants who exhibited a delayed negative EEM above the median but an immediate negative EEM below the median, where negative emotional arousal mainly lead to more efficient consolidation but did not enhance encoding (Figure 2 B). Consistent with the observed correlations between the negative and positive EEMs the immediate negative EEM group showed also a greater immediate positive 59EM (t_(213,3)_ = 5.03, p < 0.001) and a smaller delayed positive EEM (t_(223)_ = −3.66, p < 0.001) than the delayed negative EEM group.

The immediate and delayed negative EEM groups showed consistent differences in the questionnaires assessing emotional processing and related symptoms, such as rumination (Figure 3). In particular, the delayed negative EEM group reported higher trait anxiety (STAI-T, t[205.6] = 2.283, p < 0.01), a stronger tendency for dysfunctional cognitive strategies of emotion regulation in response to negative events (CERQ, t_[210.3]_ = 2.21, p = 0.03) as well as psychopathological symptoms (SCL-90; seven of nine subscales p < 0.1). By contrast, the groups did not differ with respect to age (t_[222.0]_ = −1.56, p=0.12), sex(Fisher’s exact test; odds ratio: 1.05, p = 0.88), education (χ^2^ = 25.46, p = 0.06), crystallized intelligence (t_[223]_ = −1.35, p = 0.18), fluid intelligence (t_[223]_ = 1.06, p = 0.29), working memory (t_[223]_ = 1.69, p = 0.09), attention (combined omission errors as well as reaction times; t_[194.6]_ = 0.79, p = 0.43 and t_[223]_ = 0.41, p = 0.69, respectively) and memory performance for words (immediate: t_[223]_ = 0.07, p = 0.95; delayed: t_[223]_ = 1.01, p = 0.31).

**Figure 3.**
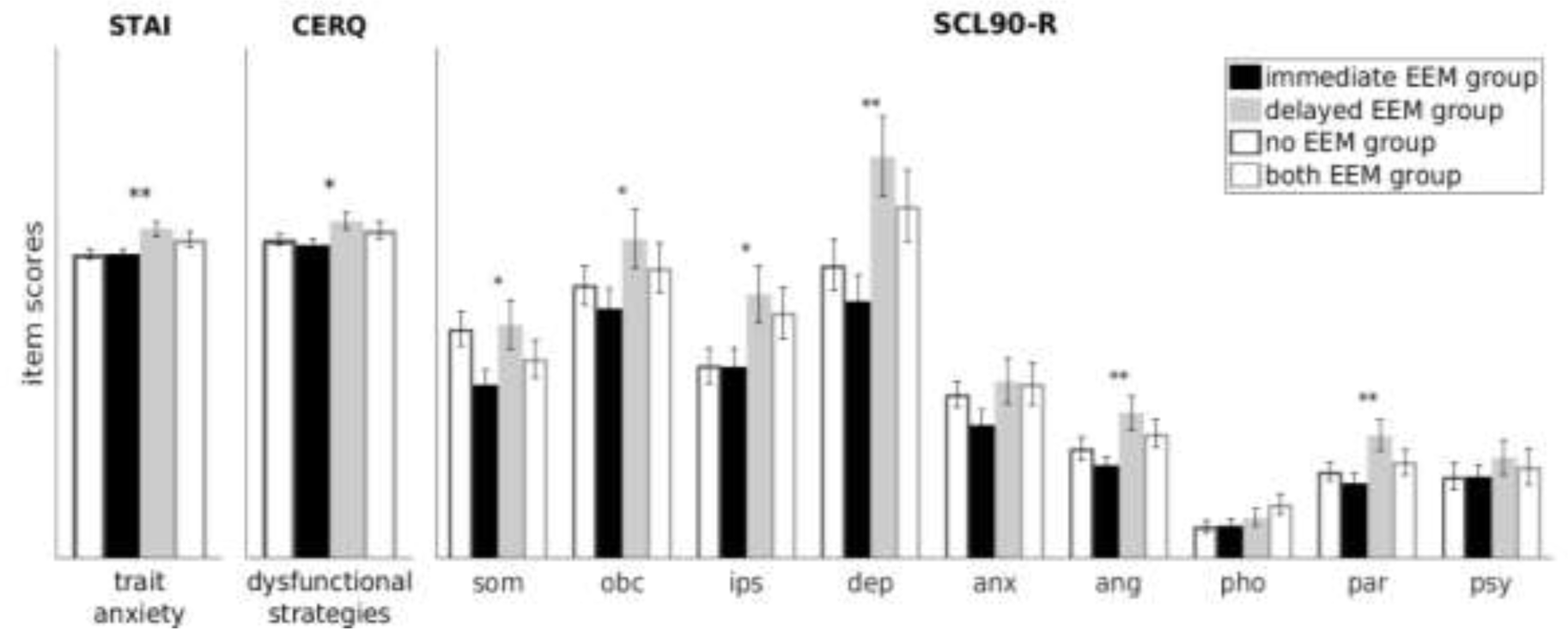
Characterization of the immediate and delayed negative EEM groups. The other two groups are shown for completeness. STAI-T, trait anxiety. CERQ, sum scores of the ‘dysfunctional’ emotion-regulation strategies (*self-blame, rumination, catastrophizing, other-blame*). SCL90-R, subscale scores (*som* = somatization, *obc* = obsessive compulsive, *ips* = interpersonal sensitivity, *dep* = depression, *anx* = anxiety, *ang* = anger-hostility, *pho* = phobic anxiety, *par* = paranoid ideation, *psy* = psychoticism). ** p < 0.01; * p < 0.05; ~ p < 0.1; mean scores and errorbars, sem.

To test whether the differences in emotional processing, between the immediate and delayed EEMs groups accounted for differences between them in negative delayed EEM we used the questionnaire scores as covariates in an ANCOVA, but this analysis still resulted in highly significant group differences (F_(1, 209)_ = 546.44, p < 0.0001). Consistent with rather subtle explanatory power of emotional processing in this analysis, a multiple regression model across the entire sample, using the delayed EEM as DV and the questionnaire scores as regressors explained only 3% of the variance (F_(11,626)_ = 1.788, p = 0.053). For completeness, we also contrasted the questionnaire scores of the corresponding extreme groups for the positive immediate and delayed EEMs, observing group differences only for the STAI (t_(230)_ = −2,38, p < 0.05, all other ps > 0.7) indicating larger trait anxiety for the delayed positive EEM group.

### 3.3 Influence of genotypes on the effects of emotional arousal on immediate and delayed memory

Finally, we explored whether the magnitude of the immediate and delayed negative EEM (corrected hit rates) depends on the genotypes in the genes coding for the α_2B_-adrenoceptor and the serotonin transporter (Figure 4). ANOVAs using the magnitude of the immediate and delayed negative EEMs as dependent variables and the α_2B_-adrenoceptor genotype as between subject factor revealed no effect of genotype (immediate negative EEM, F_[2, 627]_ = 1.85, p = 0.16, delayed negative EEM, F_[2, 627]_ = 0.72, p = 0.49). Also when the participants heterozygote for the deletion were grouped together with the participants homozygote for the deletion as done in previous publications (de Quervain et al., 2007) and the immediate and delayed negative EEMs were contrasted between carriers vs. non-carriers of the deletion there was neither a significant effect for the immediate (p = 0.22) nor delayed EEM (p = 0.36), respectively. Similarly, investigating the serotonin transporter polymorphism also showed no significant effects of genotype on the immediate (F_[2, 638]_ = 0.04, p = 0.96) or delayed negative EEM (F_[2, 638]_ = 0.39, p = 0.68). The corresponding analyses for the immediate and delayed positive EEMs also revealed no effect of genotype (all p’s > 0.40).

**Figure 4.**
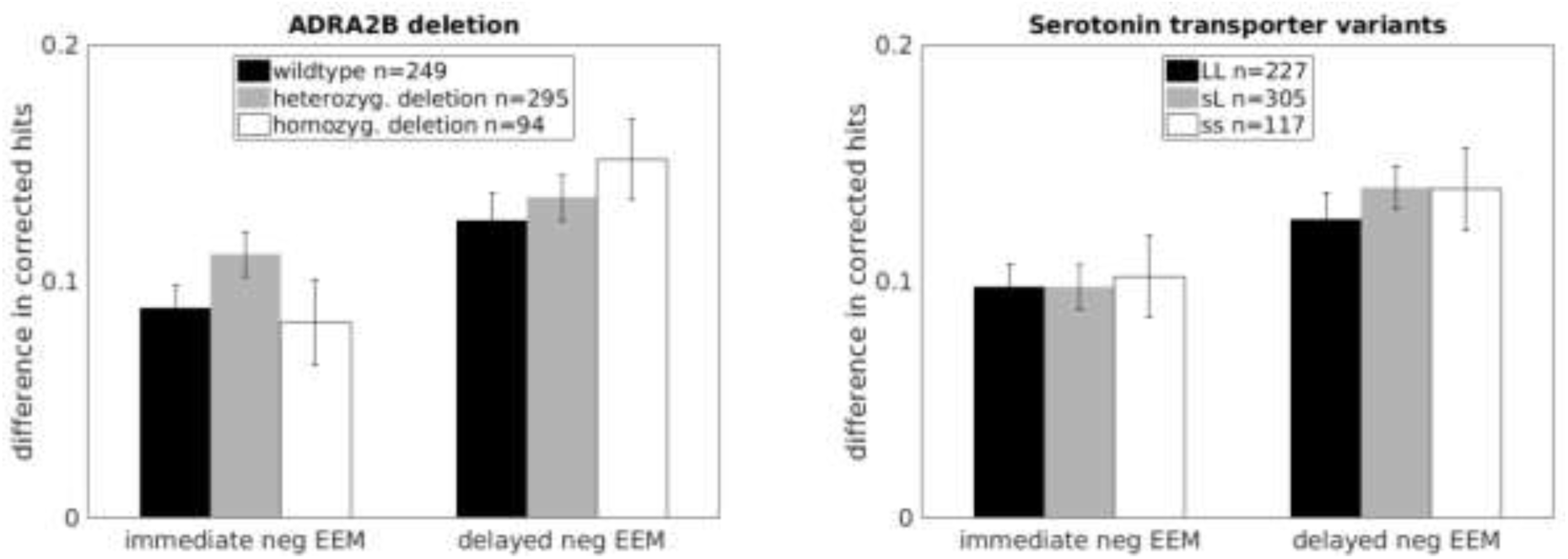
Immediate and delayed negative EEMs as a function of genotype in the genes coding for the α_2B_-adrenoceptor and the serotonin transporter.

## 4 Discussion

### 4.1 Effects of emotional arousal on immediate and dealyed memory performance

Recognition memory for emotional pictures was superior to memory for neutral pictures immediately after encoding. For familiarity and d-prime this advantage remained consistent, but for recollection, corrected hit rate, and accuracy (AUC) this advantage grew after a consolidation delay, replicating previous findings (Sharot, Verfaellie, & Yonelinas, 2007; Sharot & Yonelinas, 2008). Compared to familiarity, recollection decreased more overnight for all pictures, consistent with the notion that memories are transformed over time into more gist-like representations (Ritchey, Montchal, Yonelinas, & Ranganath, 2015; Roberts, Tsivilis, & Mayes, 2013; Sekeres et al., 2016; Viskontas, Carr, Engel, & Knowlton, 2009). Both immediate and delayed EEMs were not only of similar magnitude for both valences but the effects of positive and negative emotional arousal on immediate as well as on delayed memory were also correlated. The magnitude of both correlations suggests partly common underlying processes consistent with previous considerations (Cruciani, 2011; Hayes, Duncan, Xu, & Northoff, 2014; Mather & Sutherland, 2011). However, the response bias differed only for negative pictures, replicating previous studies (Dougal & Rotello, 2007; White et al., 2014). Importantly, compared to negative pictures, the arousal ratings for positive pictures were lower (Figure 1 A). These differences implicate that also other process than arousal and attention contributed to the positive EEMs. Consistent with this conclusion, it has been shown that the immediate EEM for positive stimuli is based also on greater semantic processing and activity in prefrontal areas whereas negative emotions more strongly enhance perceptual processing and activity in sensory areas, consistent with a larger effect on attention (Mickley Steinmetz, Addis, & Kensinger, 2010; Mickley Steinmetz & Kensinger, 2009; Talmi, Schimmack, Paterson, & Moscovitch, 2007; Yegiyan & Yonelinas, 2011). Valence-specific difference in delayed EEM have, to our knowledge, not been investigated yet. However, the positive pictures of the current study included erotic scenes and baby faces, both increase putatively dopaminergic activity in the reward system (Demos, Heatherton, & Kelley, 2012; Glocker et al., 2009); dopamine enhances memory consolidation similar to noradrenaline by synaptic tagging (Redondo & Morris, 2011; Wittmann et al., 2005). Depressive symptoms and dysfunctional emotion regulation strategies that might result in rumination and better delayed memory for the negative pictures (Kuo et al., 2012) can account only for a – albeit subtle – part of the delayed negative EEM. The lower correlation of the delayed compared to the immediate negative and positive EEMs suggests more diverging mechanisms underlying the consolidation than the encoding of negative and positive pictures.

Taken together, the pattern of results and the characteristics of the EEMs observed in the current experiment are consistent with previous findings. Moreover, the observed substantial correlations of the effects of negative and positive emotions on memory also extend previous findings by quantifying the contribution of the shared underlying processes (Mather & Sutherland, 2011). At the same time, the observation that correlations are still clearly lower than 1 together with the lower arousal elicited by the positive pictures and their rewarding nature is consistent with additional cognitive and neural processes that enhance memory only for positive stimuli. Interestingly, because the immediate correlated stronger than the delayed EEMs common processes seem to play a larger role for the immediate EEMs than for the delayed EEM.

### 4.2 Correlation of the effects of emotional arousal on immediate and delayed memory

Critically, we observed a very low correlation between the immediate and delayed EEMs across participants. This result was robust as it was not an artifact due to low reliability of the immediate and delayed EEMs, it was replicated in an independent sample, correlated substantially for negative and positive arousing stimuli, and observed moreover not only using the corrected hit rate, d-prime, familiarity and recollection estimates but also an accuracy measurement AUC that is not influence by response bias (Dougal & Rotello, 2007). Exploring the memory performance of the immediate and delayed EEM groups showed that for some participants emotional arousal enhanced only delayed but not immediate memory performance, which somehow mirrors the studies where arousal was induced only after encoding neutral items resulting only in enhanced consolidation but not immediate memory (Bass et al., 2012; Schwarze et al., 2012). This consistency suggests that emotional arousal did not result in deeper processing of the pictures for these participants. Vice versa, for other participants emotional arousal affected mainly the encoding of the pictures resulting in an immediate memory benefit that was not stable which further supports the relative independence of its effects on encoding and consolidation. Interestingly, participants who showed enhanced consolidation of negative emotional pictures, i.e. less forgetting of negative events, reported anxiety and dysfunctional emotional processing as well as more psychopathological symptoms, suggesting shared underlying neural and neuroendocrine pathways. However, alternatively participants indicating dysfunctional emotion regulation strategies and more depressive symptoms might have increased propensity to ruminate on the negative pictures which might result in reduced forgetting.

### 4.3 Neural substrates of immediately enhanced emotional memories

The pathways leading to the more efficient consolidation of (negative) emotionally arousing stimuli have been characterized intensely. In particular, a large body of animal work has shown that the activation of the central LC-NOR system and the amygdala modulate consolidation via synaptic tagging within the hippocampus (Bergado et al., 2011; McIntyre et al., 2012). The neurobiological basis of the attraction of attention by emotional stimuli has also been attributed mainly to the central NOR system and involves three main sub-processes: pre-attention/evaluation, reorienting, and sensory amplification (Carretié, 2014; Pourtois, Schettino, & Vuilleumier, 2013; Sara & Bouret, 2012). At the neural level, the potential arousal associated with a stimulus is analyzed by prefrontal and subcortical areas, most importantly the amygdala (Pessoa and Adolphs 2010). According to the most prominent view, the prefrontal cortex and central amygdala then recruit the central NOR system by activating the LC (Markovic, Anderson, & Todd, 2014; Mather, Clewett, Sakaki, & Harley, 2015; Sara & Bouret, 2012). NOR drives the ventral attentional network to “reset” the dorsal top-down network and to promote the shift in attention (Sara & Bouret, 2012). Moreover, the LC re-projects NOR to the BLA, and also to the hippocampus, where NOR regulates synaptic plasticity (Hagena, Hansen, & Manahan-Vaughan, 2016; O’Dell, Connor, Guglietta, & Nguyen, 2015). In addition, the LC projects directly to all cortical areas, including the visual processing stream, which enhances activity and improves signal-to-noise ratio (Edmiston et al., 2013; Mather et al., 2015). The enhanced attention allocated to emotional stimuli, a result of the activation of the central LC-NOR system, finally results in a greater probability of successful hippocampal encoding. The activity in prefrontal, parietal and visual areas that is observed in fMRI-studies during successful encoding of arousing stimuli is consistent with reflecting these attentional processes (Dolcos et al., 2012; Murty et al., 2010; Talmi et al., 2008).

### 4.4 Dissociable noradrenergic pathways underlying effects of emotional arousal on immediate and delayed memory

On its own, it is therefore possible that the same mechanisms (activation of the central LC-NOR system) that enhance immediate memory will also give rise to the preferential consolidation of prioritized traces. However, the low correlation of both effects in the current study suggests at least partly distinct neurobiological substrates. One explanation for this observation would be that activity in visual processing areas leading to more successful encoding and better immediate memory for emotional items is upregulated not by the central LC-NOR system but directly by the amygdala as a response to arousal detection (Chen, Li, Jin, Shou, & Yu, 2014). However, lesion studies suggest that emotional biases of attention do not require an intact amygdala, implying a critical role for the central LC-NOR system (Bach, Talmi, Hurlemann, Patin, & Dolan, 2011; Piech et al., 2011). Therefore, our findings support an alternative hypothesis, namely that processes downstream of the activation of the central LC-NOR system have different effects on encoding and consolidation. Such functional differentiation is likely because this system exhibits a marked anatomical heterogeneity. Distinct populations of LC neurons project to the amygdala and cortex, on the one hand, and to the hippocampus, on the other (Schwarz & Luo, 2015). In addition, most LC-NOR neurons do not form one-to-one contacts with postsynaptic neurons but have a more diffuse hormone-like action in the brain. The proportion of one-to-one contacts differs between brain regions, e.g. about 15% in the hippocampus, only 5% in cortex and up to 45% in the amygdala (Vizi, Fekete, Karoly, & Mike, 2010; Zhang, Muller, & McDonald, 2013). These anatomical differentiations in terms of projection targets and modulation specificity could be a basis for the observed functional differentiation of the effects of an LC-NOR activation. For instance direct LC-NOR projections to the hippocampus and visual cortex might modulate encoding as a response to enhanced attention without involving the amygdala - consistent with the lesion data on emotional biased attention (Bach et al., 2011; Piech et al., 2011). Contrary, the critical role of the amygdala in modulating hippocampal activity after LC-NOR activation is well described (Roozendaal & McGaugh, 2011). Most importantly, in the hippocampus distinct molecular pathways might be involved in facilitation of early LTPs/LTDs (immediate EEM) and synaptic tagging (delayed EEM), both triggered by a phasic increase in NOR (Hagena et al., 2016; O’Dell et al., 2015).

A complementary neurobiological basis for differential effects of an LC-NOR activation could be the differential involvement of the adrenoceptor families or subtypes in both effects because they differ with respect to their affinity and action (Hein, 2006). The role of β-and α_1_-adrenoceptors in consolidation has been shown in many experiments (Roozendaal & McGaugh, 2011). The role of the α_2_-adrenoceptor family has been studied using the polymorphism in the gene coding for the α_2B_-adrenoceptor (Todd et al., 2011). This polymorphism might be related to enhanced amygdala activity, subjective vividness, and attraction of selective attention during emotional processing (Rasch et al., 2009; Todd et al., 2013, 2015). It was also observed that the α2B-adrenoceptor polymorphism affects the immediate EEM in free recall (de Quervain et al., 2007; Rasch et al., 2009). However, the immediate EEM in free recall disappears when one controls for the higher distinctiveness and semantic relatedness of emotional stimuli (Barnacle, Montaldi, Talmi, & Sommer, 2016), questioning which process was influenced by the polymorphism in these studies. We did not observe an association of this polymorphism with the immediate EEM in recognition tests but also did not observe an association with the effect on consolidation. Therefore our results do not support the hypothesis of a differential involvement of the adrenoceptor in the effect of emotional arousal on encoding and consolidation. Finally, also the serotonin-transporter genotype did not correlate with neither the immediate nor delayed EEM.

## 5 Conclusion

In conclusion, our data show an intriguing dissociation of the effects of emotional arousal on immediate and delayed recognition memory. Previous work has established that the better immediate recognition of emotional stimuli is based on the allocation of more attention to emotional stimuli, and that the LC-NOR system would have been activated. The enhanced delayed memory for emotional stimuli is due to their more efficient consolidation which depends also on the LC-NOR system. The low correlation between immediate and delayed EEM – our central finding – must, therefore, reflect partly independent effects of NOR on downstream neural processes. We propose that individual differences in the effects of NOR on cortical signal to noise ratio and early hippocampal LTP, on the one hand, and their conversion into late LTPs, on the other, underlies individual differences in immediate and delayed EEM.

## Acknowlegements

This work was supported by a State of Hamburg excellence initiative (Landesexzellenzcluster 12/09 “neurodapt”).

